# Gearing up Golden Braid assembly for plant synthetic genomics with RepTiles

**DOI:** 10.1101/2025.02.28.640145

**Authors:** Viktoria Petrova, Dejan Andrejic, Tobias Finkenrath, Josepha Grewer, Matias D. Zurbriggen, Uriel Urquiza-Garcia

**Affiliations:** Institute of synthetic Biology, Düsseldorf University, Düsseldorf, Germany; Cluster of Excellence on Plant Sciences CEPLAS, Germany

**Keywords:** plant synthetic genomics, biodesign automation, phytobricks, golden gate, random DNA, golden braid, synthetic biology

## Abstract

We have developed RepTiles, a system for generating traces of random DNA for synthetic genomics applications. RepTiles reduced the burden associated with the manual design of long DNA from standard biological parts. We are applying RepTiles to the construction of random DNA segments that will form part of a neochromosome in *Physcomitrium patens*. RepTiles has a base DNA collection of 52 1.7 kb chemically synthesised Phytobricks mini-chunks. We provide a user-friendly web application that facilitates the design of DNA assemblies based on the base collection. The system generates assembly plans for generating chunks using Golden Braid. The resulting chunks can then be assembled into megachunks using Transformation Associated Recombination cloning. It thus supports the generation of ultra-long synthetic DNA from phytobricks, contributing to the vision of synthetic plant genomics from modular parts to complete genomes.

## Introduction

Synthetic genomics has progressed from the construction of small bacteriophage genomes to the synthesis of artificial yeast genomes (Venter et al., 2022). Plant genomes represent the next frontier in synthetic genomics. The latter has been initiated in *Physcomitrium patens (P. patens)* with the re-coding of ∼155 kb of the short arm of chromosome 18 into ∼68 kb designer DNA (Chen et al., 2024). The plant synthetic biology community has been creating and characterising biological parts that follow the Phytobricks standard. However, there is no current path forward on how to go from standard biological parts into full genomes. RepTiles 1.0 starts to fill this gap. In particular in our efforts to build a fully synthetic neochromosome in *P. patens*. Random DNA chromosomes, with a 18 kb length have been used in yeast, and functionally characterised transcriptionally and epigenetically (Gvozdenov et al., 2023). We see the potential for such neochromosome for providing valuable information about chromosome biology, genomic level functional studies or adding new functionality to the *P. patens* genome. For example, users might be interested in the introduction of large metabolic pathways for the production of high-value compounds like taxol (Liu et al., 2024).

We are approaching the construction of this neochromosome with a modular and standardized approach, using the Golden Braid assembly standard (Vazquez-Vilar et al., 2017). As a first step, we have developed RepTiles, an extension of Golden Braid for creating long tracks of random DNA from Phytobricks (Patron et al., 2015; Vazquez-Vilar et al., 2017). The platform includes software to automate the design of assembly plans based on Golden Braid and Transformation Associated Recombination (TAR) cloning (Kouprina et al., 2021). This two-step approach starts with the expansion of standard mini-chunks into DNA chunks using the iterative and hierarchical properties of GB assembly. In the second step, mid or mega-chunks of DNA can be assembled from the chunks using TAR cloning, thanks to the overlapping nature of the chunks designed by RepTiles. The platform has a base set of 52 mini-chunks of random DNA domesticated in the GB universal domestication plasmid pUPD2. The basic mini-chunks provide the raw material for the generation of random DNA fragments up to 90 kb. We tested the fragment expansion by generating two pieces of ∼14 kb, validating our approach.

This bioware and software provides valuable material for synthetic plant genomics, in particular for the creation of reusable plant artificial chromosomes (rPACs). The strategy presented in this work using automation and balancing between Golden Braid and TAR cloning provides a way to generate neutral genetic material onto which biological functionality can be loaded by homologous recombination in *P. patens*. In addition, established neutral loci can be further extended by random tracts of DNA, providing new neutral recombinogenic regions. Therefore, we see an important potential to generate large stretches of random DNA for plant synthetic genomics applications. It is now possible to explore the use of random DNA thanks to the significant decrease in the cost of chemically synthesised dsDNA.

## Results

### RepTiles, a tool for automating the extension of random DNA fragments for synthetic genomics applications

The RepTiles assembly tool is a Flask(Python)-based web application designed to scale up the construction of large constructs from standard PhytoBricks (see supporting information). RepTiles provides the logic for GB assembly of overlapping chunks that can be assembled into mid- or mega-chunks using TAR cloning. RepTiles is containerised using Docker and can be launched using docker-compose up. This allows it to be quickly deployed across different operating systems and gives users full control over their data flow. RepTiles uses the open source cloning simulation framework DNA Cauldron for *in-silico* verification and visualisation of the assembly design (Edinburgh Genome Foundry, University of Edinburgh, Edinburgh, UK). The latter predicts the final construct sequences and identifies potential assembly errors. This is proving useful in assisting users by simulating the maps used in the validation of large-scale, multi-step DNA assembly processes. Map generation is a very attractive feature of DNA Cauldron and we decided to reuse it. This was made possible by the open source nature of DNA Cauldron. This automation reduces the chance of human error in the manual generation of plasmid maps and also reduces the workload on the human side.

RepTiles is complemented by a collection of domesticated random DNA fragments in the pUPD2 universal domestication vector of GB (Supplementary Table 1). The user has to enter the Genbank files (.gb) for the collection of GB assembly plasmids (3α1, 3α2, 3Ω1 and 3Ω2), the collection of Phytobricks mini-chunks domesticated in pUPD2 with overhangs A1-C1, the number of chunks and the length of each of them. With this information, the software designs an assembly plan to generate each of the chunks, stacked as close as possible to the length specified by the user. The generated assembly plans can be downloaded directly by the user or fed into an internal instance of DNA Cauldron. This simulates the assemblies based on the assembly plans calculated by RepTiles and runs asynchronously using Celery and Redis. When the simulation is complete, the user can download the DNA Cauldron results as a zip file. This file contains the collection of .gb files for each of the intermediate steps during chunk construction, which can be used as a reference for verifying each chunk. The generated chunks are designed so that one DNA chunk ends with a particular mini-chunk sequence, and the same sequence is used in the first part of the next chunk. The chunks are designed by concatenating the sequences of the input fragments until they reach the maximum user-defined length. Fragment names and their respective sequences are recorded in tuples for all chunks. In this way, RepTiles returns a list of chunk assemblies. The finished chunks can be used for mega-chunk construction by TAR cloning thanks to the presence or recombination arms.

Due to the hierarchical and iterative nature of the Golden Braid, the assembly of each chunk can be performed in parallel. To represent this structure of hierarchical assemblies, the RepTiles contain a Python class (GB_fragment) with the necessary attributes and associated functions (Supplementary Table 4 and Table 5). Each instance of the class represents an input fragment with attributes such as order, size, fragment ID (essentially the name of the fragment .gb file). An assembly of two fragments (“parents”) results in a new class instance (“child”), which inherits the sequences of the parents (assemble function). In addition, the child’s attribute origin is set to the parents’ instances and each parents’ attribute destination is set to the child, resulting in a tree structure.

Initially, RepTiles builds an “empty” tree structure by simply combining two objects to create a new one, until there is only one object (Figure 1). The initial set of fragments to be concatenated into a tree is determined by the find_and_concatenate_sequences function. Once the structure is established, a backpropagation would distribute a specific Golden Braid backbone to each object. This is recorded by the class instance “level”, which is a numpy array. The propagation starts at the top of the tree, which is the last object from the assembly. By default this object would have an alpha 1 backbone ([0, 0]). The algorithm then looks for the origin of this object (the parents) and assigns the appropriate backbones based on the Golden Braid assembly standard (in this case omega 1 and omega 2). More specifically, the algorithm checks the first value of the child’s level array (0 for alpha, 1 for omega) and then distributes the opposite value to the parents. One of the parents gets a 0 for a second array value, while the other gets a 1 to account for compatibility of sticky ends (alpha 1 and 2; omega 1 and 2). In this way, level instances are distributed to all objects in the tree. This assembly standard allows parts to be assembled in a predictable way, as the backbone determines where the insert would end up in a next round of assembly. Thus, backpropagation also provides information about the order of the fragments in the final construct. This can be further exploited by users if they wish to use existing GB parts that have already been cloned into alpha or omega backbones.

**Figure 1.**
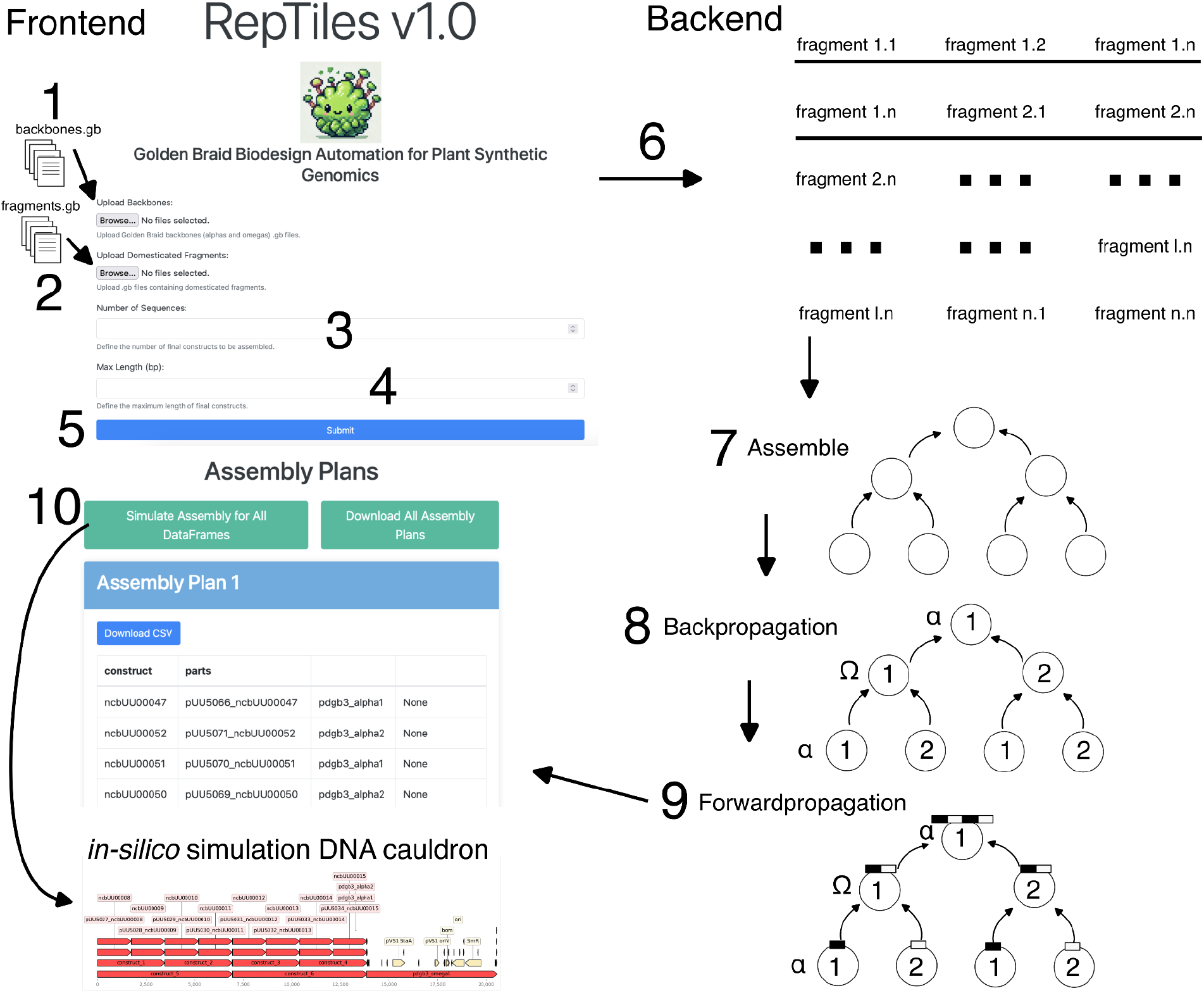
RepTiles frontend and backend. (1) The RepTiles frontend takes as input Golden Braid backbones and the level 0 fragments to be assembled (2). (3) The user defines the number of chunks to be assembled and their size in bp (4) and can submit (5) for design. The backend generates the assembly plan, which is then taken by the assemble function (7) that creates a tree that specifies the location with the backpropagation function (8). This tree is then filled up with the forwardpropagation function (9) returning the assembly plans. These assembly plans can be downloaded by the user or fed to RepTiles instance of DNA Cauldron, which generates assembly reports. Among the output files, the user receives Genbank files for each assembly step, which are used for assembly verification (10). Circles represent alpha or omega backbones from Golden Braid.

**Figure 2.**
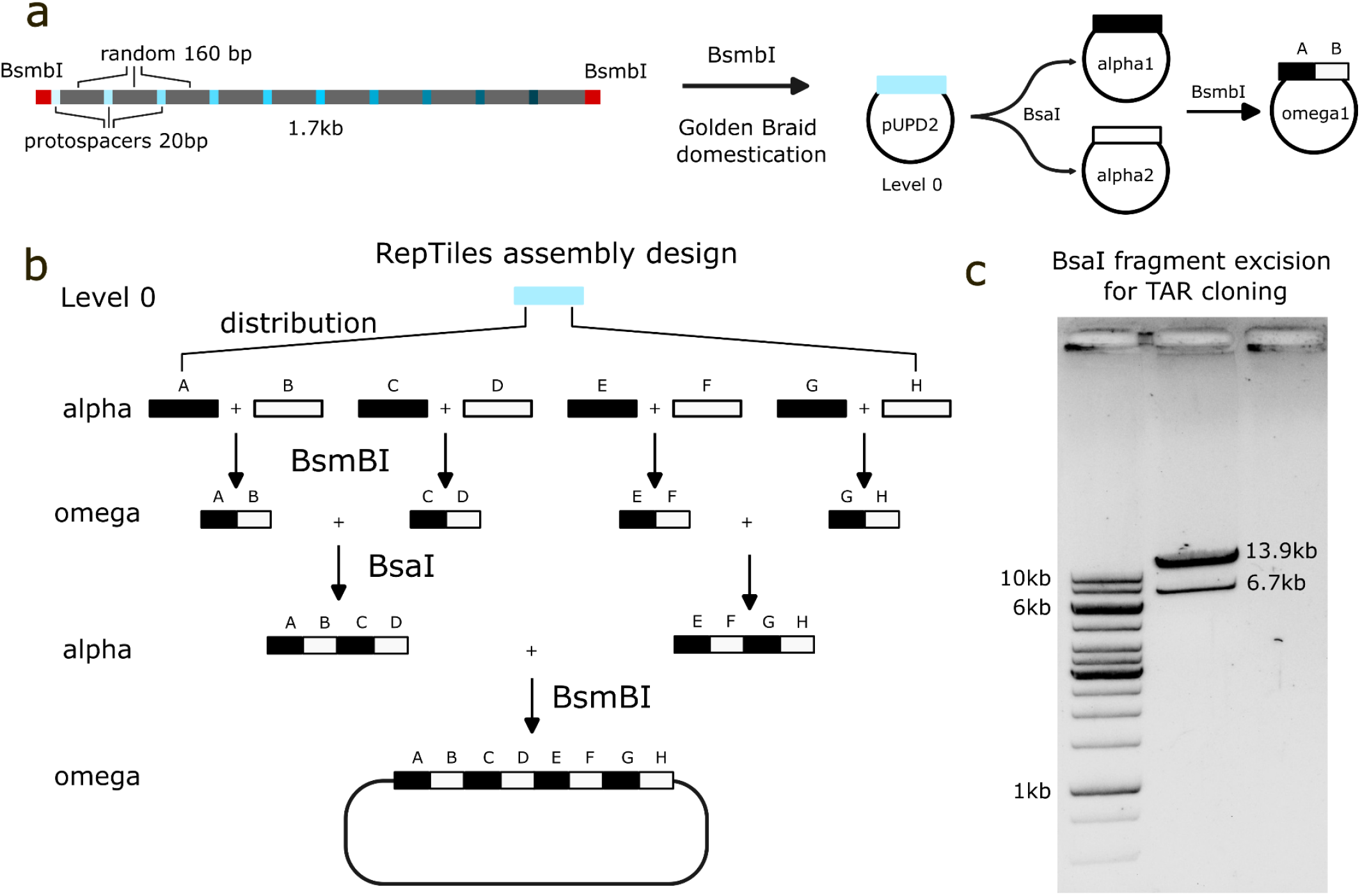
Golden Braid assembly of a ∼14 kb random DNA track designed with the aid of RepTiles. a) Design and domestication of random DNA mini-chunks. Direct domestication from synthesis into Golden Braid universal domestication plasmids pUPD2. Level 0 parts can be distributed to position 1 or 2 on the alpha level. These parts can then be further used and cloned into an omega vector. b) RepTiles automates the assembly design using the Golden Braid mechanisms until we reach an omega level that can be easily excised with BsaI for further assembly. c) Digestion of a synthetic DNA track constructed using RepTiles with BsaI, which can be further used for yeast TAR cloning.

As mentioned above, backbone distribution occurs during backpropagation, starting from the top of the assembly tree. By default, the first backbone is always a pDGB3α1. However, when the empty tree structure is created, or just before backpropagation starts, the probability of having more alpha backbones than omegas in the first level is calculated (calculate_alpha_probability function). If this probability is less than 0.5, the default backbone at the top is changed to pDGB3Ω1. This function was introduced because when parts are to be moved from level 0 to an omega vector, the reaction involves the use of two different Type-IIS restriction enzymes (BsmbI and BtgZI). BtgZI releases parts from the domesticator backbone, while BsmBI cleaves the receiving omega backbone. In contrast, the reaction to move from pUPD2 backbone to an alpha backbone requires only one restriction enzyme - BsaI. As a result, the latter is preferred because it is more efficient and cost-effective. Thus a function to optimize the number of alpha backbones used in the first stage of assembly. It generates the assembly plan in such a way that its execution in the laboratory requires the use of fewer enzymes, thus reducing assembly costs. This is particularly important in the context of synthetic genomics projects.

### The design of the standard RepTiles random DNA library and the assembly of two ∼14 kb chunks

As part of RepTiles, we provide a collection of 52 mini-chunks of 1.7 kb random DNA that can theoretically be assembled into a 90 kb fragment based on RepTiles. The mini-chunks lack the presence of BsmBI, BsaI and BtgZI restriction sites. To each 1.7 kb sequence, we have added 10 sequences of 20 bp length with an upstream PAM of 150 bp spacing. The user can target this random fragment at once using CRISPR/Cas9-derived technologies. They also contain adapters for direct Golden Gate domestication into pUPD2. This design allows the complete assembly to be executed without PCR. Using the RepTiles collection, we designed a ∼90 kb mega-chunk. We tested the mechanics of GB assembly on two ∼14 kb chunks with successful results verified by Oxford Nanopore full plasmid sequencing.

## Discussion

We started to close the gap between standard Phytobrick parts and synthetic genomics using random DNA as part of our construction efforts of a synthetic neochromosome in *Physcomitrium patens*. The potentially large scale requirements for stable plant chromosomes requires the use of automated tools for biodesign. RepTiles provides a convenient tool for automating the design of large random synthetic constructs based on the Golden Braid assembly standard and TAR cloning. We have implemented RepTiles using a containerised approach to facilitate portability between platforms. In addition, RepTiles provide users with full control over the flow of data, reducing the potential for compromise when submitted to publicly hosted web services. We tested the mechanics of RepTiles by successfully synthesising two ∼14 kb pieces of DNA as part of the synthesis of a random *P. patens* neochromosome.

RepTiles complements other tools that are emerging, such as GenoDesigner, which was designed to facilitate the re-coding of the *P. patens* genome as part of the SynMOSS project (Yu et al., 2024). GenoDesigner does not provide the support needed to generate tracts of random DNA from phytobricks. RepTiles helps us to fill this gap. Once a synthetic random neochromosome is in place, GenoDesigner will be useful for adding functionality to the random DNA chromosome. In the future, incorporating features such as multi-part assembly logic and enhanced user control over fragment arrangement would enhance RepTiles’ capabilities and ensure that it continues to meet the evolving needs of large-scale synthetic biology projects. By taking inspiration from the success of other tools, RepTiles can continue to evolve as an indispensable asset in the rapidly emerging field of plant synthetic genomics.

## Methods

### Fragment synthesis and domestication

Fragments were generated automatically using a Python script that generated random samples of A, T, G, C with equal probability. A 20 bp plus NGG PAM was added every 150 bp for future dCas9 targeting. Sequences containing any of the following enzymes: BsmbI, BtgZI, BsaI or BpiI were resampled. Domestication adapters were added at the end of each sequence to allow domestication of BsmBI into the pUPD2 backbone. The position of the domesticated random DNA spans from position A1-C1, allowing it to be directly transferred to the alpha or omega acceptor backbone. The synthetic mini-chunks were ordered from Twist Bioscience and cloned directly from synthesis into pUPD2 using BsmBI Golden Gate cloning. Fragments were resuspended in TAE to an estimated concentration of 75 ng/ul and quantified using NanoDrop. A Golden Gate domestication reaction was performed with 1 ul of synthetic DNA and 75 ng/ul of pUPD2, 1 ul of 10 units of BsmBI (New England Biolabs), 1 ul of DTT 10 mM, 1 ul of ∼3 units of T4 ligase (New England Biolabs). and 1 ul of T4 ligase buffer (New England Biolabs). The reaction was incubated for 25 cycles between 5 min at 16ºC and 5 min at 45ºC and the enzymes were inactivated 10 min at 60ºC and then 10 min at 80ºC. 2 ul of the reaction was then transformed into chemically competent TOP10 (Thermo Fisher Scientific) or NEB 10-beta (New England Biolabs) cells. Transformants selected on LB supplemented with 35 ug/ml Cm, 0.5 mM IPTG, 0.5 mM and 40 ug/ml X-Gal followed by colony PCR. Candidates were then verified by restriction digestion and stored at -20ºC until further use.

### Golden Braid fragment expansion

The hierarchical assembly of the ∼14 kb chunk number 2 was performed via a series of Golden Braid reactions (see Supplementary Table 4 for the list of plasmids). The established general workflow was as follows: GB assembly reaction, transformation of chemically competent NEB 10-beta E. coli cells (New England Biolabs), colony PCR, mini-plasmid preparation and restriction test digestion. The GB assembly reactions are summarised in the table below:

| GB alpha/omega |  |
| --- | --- |
| Insert 1 | 75 ng |
| Insert 2 | 75 ng |
| alpha/omega backbone | 75 ng |
| BsaI 20 U/ul (NEB)/ BsmBI 10 U/ul (NEB) | 0.5 ul |
| T4 ligase 4,000 U (NEB), diluted 1:1000 | 1 ul |
| If BsmBI use 10 mM DTT | 1 ul |
| T4 ligase buffer (NEB) | 1 ul |
| ddH<sub>2</sub>O | Up to 10 ul |

**Alpha** reactions are incubated 25 cycles of 2 min at 37ºC and 5 min at 16ºC then 10 min at 60ºC followed by 10 min at 80ºC. **Omega** reactions are incubated 25 cycles of 5 min at 45ºC and 5 min at 16ºC then 10 min at 60ºC followed by 10 min at 80ºC. Then 2 ul of the mixture is transformed into NEB 10-beta *E. coli*.

### Full plasmid sequencing

The correct assembly of the two ∼14 kb random DNA fragments (pUU5156 and pUU5194) was assessed by Oxford Nanopore Technologies full plasmid sequencing, which was performed by Microsynth AG Göttingen. Prior to sending samples for sequencing, 2 replicates were prepared with a concentration of 100 ng/μL according to the NanoDrop. In addition, a Qubit working solution was prepared by mixing 1 x n μL Qubit™ reagent with 199 x n μL Qubit™ High Sensitivity Buffer, where n = 4 (2 replicates + 2 standards). 10 μL of the two standards supplied with the kit and 1 μL of the sample were mixed with 190 μL and 199 μL of the working solution, respectively. The standards were measured first, followed by the samples. The average of the replicates was taken as the final concentration. The sample was sent for sequencing in a final volume of 15 μl with a concentration of 20 ng/μl. Results were analysed by aligning the sequence to the plasmid map automatically generated by RepTiles in Benchling [Biology Software]. (2025), retrieved from https://benchling.com.

## Author contributions

V.P., M.D.Z and U.U.G designed the study. V.P. and U.U.G. designed the software architecture. V.P. wrote the code with input from U.U.G. V.P., D.A., T.F. and U.U.G performed the experiments. J.G. and U.U.G. supervised the experimental work. M.D.Z. and U.U.G. acquired funding and supervised the work. V.P. and U.U.G. drafted the first version of the manuscript. All authors contributed to and approved the final version of the manuscript.

## Acknowledgements

We thank Leonie-Alexa Koch for contributing to the naming of RepTiles. This study was funded by the Deutsche Forschungsgemeinschaft (DFG, German Research Foundation) under Germany’s Excellence Strategy – EXC-2048/1 – project ID 390686111 to J.G., M.D.Z. and U.U.G. In particular, with a seed fund awarded to U.U.G. by the Cluster of Excellence on Plant Sciences (CEPLAS).

## Supporting information

The code can be accessed in the public repository https://github.com/jurquiza/reptiles instructions can be followed for installation using Docker and docker compose.

